# Neutral lipids restrict the mobility of broken DNA molecules during comet assays

**DOI:** 10.1101/2024.10.11.617826

**Authors:** Caroline Soulet, Jordi Josa-Castro, María Moriel-Carretero

**Affiliations:** Centre de Recherche en Biologie cellulaire de Montpellier (CRBM), Université de Montpellier, Centre National de la Recherche Scientifique, 34293 Montpellier cedex 5, France; Institut de Génétique Moléculaire de Montpellier (IGMM), Université de Montpellier, Centre National de la Recherche Scientifique, 34293 Montpellier cedex 5, France

**Keywords:** comet assay, neutral lipids, broken DNA, acetone

## Abstract

One widespread technique to assess in relative terms the amount of broken DNA present in the genome of individual cells consists in immobilizing the cell’s nucleus under an agarose pad (called the nucleoid) and subject the whole genome to electrophoresis to force broken DNA molecules out of it. Since the migrating DNA molecules create a tail behind the nucleoid, this technique is named the comet assay. While performing comet assays regularly, we systematically observed circular regions devoid of DNA within the nucleoid region. We characterize here that these correspond to clusters of neutral (apolar) lipids, since they could be labelled with neutral lipid-dying molecules, increased when cells were fed with oleic acid, and were irresponsive to the electrophoretic field. Of relevance, de-lipidation assays either *in vivo*, or *in vitro*, show that these neutral lipids within the nucleoid limit the ability of broken DNA molecules to migrate into the comet tail. From a technical point of view, we show that de-lipidation permits a wider range for the detection of broken DNA molecules. Biologically, we put forward the notion that neutral lipids in contact with DNA may locally exert regulatory functions within the cell’s nucleus.

## Introduction

Genome integrity is put at risk by multiple challenges, both from endogenous and exogenous origin, that harm the DNA molecule in different ways. One very toxic DNA damage is its break at either one or both strands, which by definition disrupts the continuity of the genetic information. Indirect markers are frequently used to establish the presence or absence of broken DNA molecules in a genome, for example the detection of a phosphorylated form of the histone H2A, γH2AX, as this mark is deposited on the chromatin surrounding these lesions (Sharma *et al*, 2012). However, if the process of signaling that drives the inclusion of this mark is hijacked, for example due to mutations or pathological scenarios, the conclusions about the underlying level of damaged DNA may be incorrect. This is why techniques directly assessing the actual physical integrity of the genome are a more reliable source of information. For example, pulsed field gel electrophoresis permits the purification of whole chromosomes (obtained from thousands of cells) into agarose plugs that are migrated in an electrophoretic field. The broken molecules migrate readily in the gel, while the intact chromosomes remain in the well or poorly enter the agarose matrix (Kawashima *et al*, 2017). A more refined technique, termed Single Cell Gel Electrophoresis, permits the assessment of broken DNA molecules at the single cell level. This technique immobilizes each cell’s nucleus under an agarose pad (called the nucleoid). Chemical treatment to remove the DNA-surrounding nuclear envelope then allows, upon being subjected to electrophoresis, to force broken DNA molecules out of the nucleoid. The migrating DNA molecules create a tail behind the nucleoid, which justifies the technique is more commonly named “comet assay” (Figure 1A) (Liao *et al*, 2009).

**Figure 1.**
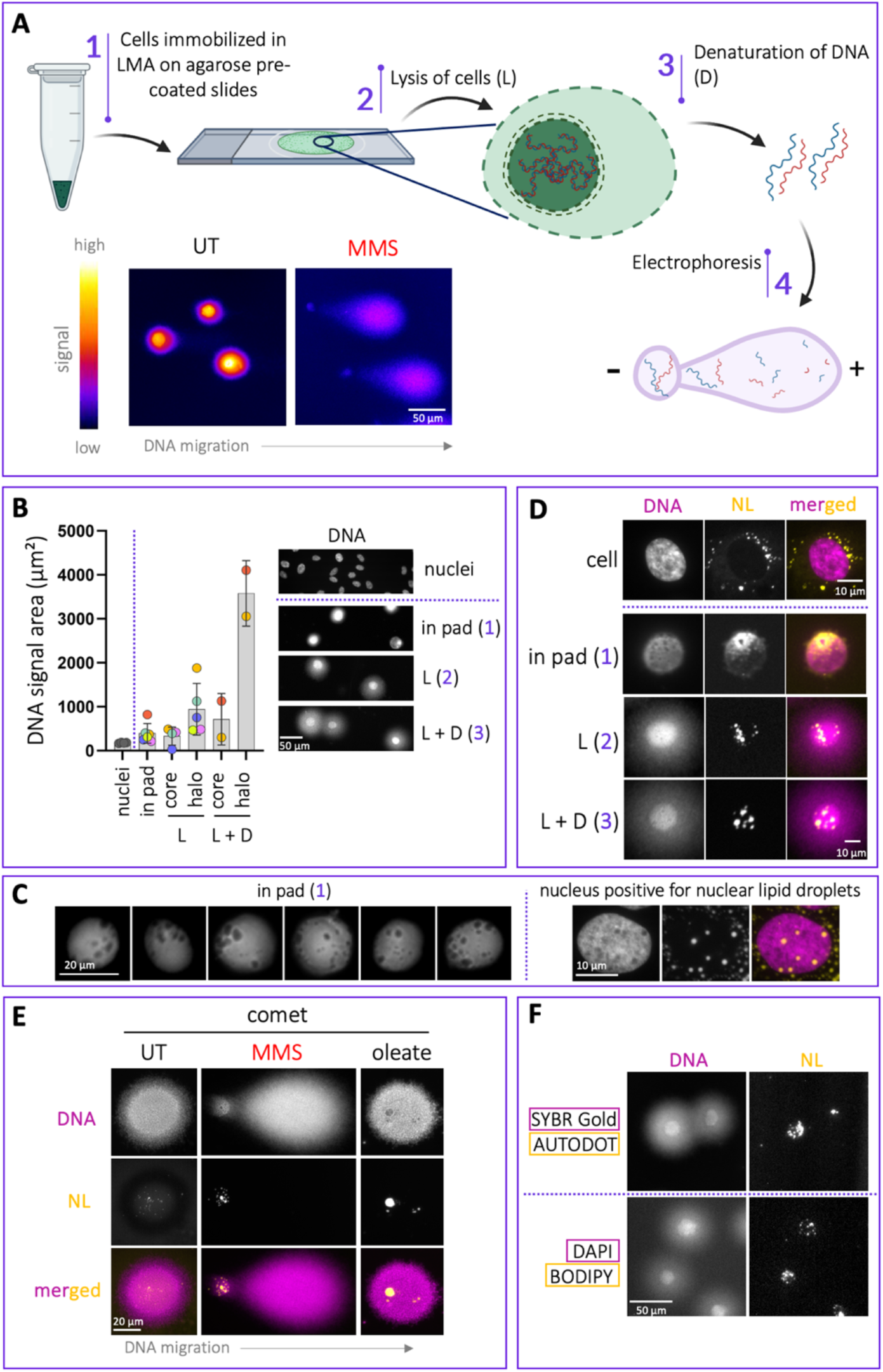
Neutral lipids (NL) invade the DNA mass during the execution of the comet assay protocol. **A**. Scheme of the protocol steps and illustration (partly created using BioRender) with real microscopy images obtained at the end of those steps by dying DNA with SYBR Gold^TM^. The DNA stems from untreated (UT) or MMS-treated (methyl methanesulfonate, leads to DNA breaks) cells. **B**. The area occupied by the DNA in intact cells (nuclei), or after execution of the protocol (halted at each individual step: in pad; after L, lysis; after denaturation, D) was measured and is plotted on the left. For L and D, the signal is subdivided in core-only, or including the halo. The dots on the graph correspond to the mean DNA area out of at least between 75 and 500 DNA objects per experiment, where each color refers to an independent experiment. The error bars represent the standard error of the mean. **C. Left:** detail of 6 different in-pad nuclei displaying “holes” devoid of DNA-dying signals. **Right:** example of one nucleus from a culture of Huh-7 cells incubated with oleate, a treatment known to elicit the formation of nuclear lipid droplets. The position of nuclear lipid droplets coincides with “holes” devoid of signal in the DNA channel. **D**. DNA was dyed as in (B), but NL were simultaneously dyed using AUTODOT^TM^. **E**. Details as in (B, D), but the cells used to perform the comet assay were pre-treated with MMS or with oleate. **F**. DNA and NL were labeled with different reagents to assess the generality of the observed phenomenon.

We regularly perform comet assays in view of our interest in understanding the mechanisms ensuring genome stability. We realized that nucleoids, visualized by DNA dyes, were not uniform in signal but contained “empty” circular regions. We are not the first ones to detect these entities, as a fast, informal query of the net showed others have wondered before what they were, and that much speculation was sparked by these reports (https://www.researchgate.net/post/COMET_assay_in_neutral_condition_Why_do_I_observe_black_dot_in_nucleus).

In this brief article, we report our characterization of these structures. We characterize that neutral lipids originally present in the cytoplasm of the cells get entangled within the DNA of the nucleoid as a consequence of the comet assay protocol. De-lipidation assays either *in vivo*, or *in vitro*, show that these lipids within the nucleoid limit the ability of broken DNA molecules to migrate into the comet tail. Technically speaking, this means that de-lipidation will permits a wider range for the detection of broken DNA molecules. On a more biological consideration, we add to the incipient body of evidence indicating that neutral lipids can restrict the mobility of DNA molecules.

## Material & Methods

### Reagents

4-NQO (Merck, N8141-250MG),

Acetone (Carlo Erba, 400971)

AUTODOT™ Visualization Dye (Clinisciences, SM1000a),

BODIPY (Merck, 790389),

Camptothecin (CPT, C9911, Sigma-Aldrich),

DAPI (D9542, Sigma-Aldrich),

Hydroxyurea (HU, Merck, H8627-25G)

Low Melting Agarose (LMA, Merck, A9414-10G),

Methyl MethaneSulfonate (MMS, 129925, Sigma-Aldrich),

Neocarzinostatin (NCZ, N9162-100UG, Sigma-Aldrich),

Oleic Acid-Albumin from bovine serum (OA, O3008, Sigma-Aldrich),

SYBR™ Gold Nucleic Acid Gel Stain 10,000X (10358492, Fisher Scientific),

Zeocin (R25001, ThermoFisher)

### Cell culture and treatments

The RPE-1 (and the Huh-7, only used in Figure 1C) cell lines used in this study were authenticated by ATCC STR profiling. They were grown in DMEM+GlutaMAX (Gibco, 31966047) supplemented with 10% FBS (F7524-500ML, Sigma) and 1% Pen/Strep (P0781, Sigma-Aldrich) at 37°C and 5% CO_2_. DNA-damaging treatments were applied 2 hours before harvesting, the final concentrations were: 4-NQO: 2.5μM, CPT: 1μM, HU: 1mM, MMS: 0.005% (or as indicated in the legend), NCZ: 120 ng/mL, and zeocin: 10 μg/mL. Loading with oleic acid was done 24h before harvesting at 60 μM. For starvation experiments, complete medium was removed from the well 20h before harvesting, and replaced by FBS-less pre-warmed medium.

### Alkaline Comet Assay

Cells were seeded on 24 well-plates (dilution 1:5 from one confluent p100), and grown for 24 hours. Cells were treated as specified, rinsed with 1X PBS then quickly trypsinized. Cell pellet was resuspended in 1X PBS to obtain a cell suspension at 10^5^ cells/mL. For controls *in vitro*, H_2_O_2_ was added at 100 μM and incubated at 4°C during 20 min. 30 μL from this suspension was mixed with 250 μL of molten 1% Low Melting Agarose (LMA), both in 1X PBS, and then 30 μL of this mix poured onto 1% agarose pre-coated coverglass, and directly covered by a 12 mm-diameter coverslip. After 10 min of polymerization at 4°C, the coverslip is removed and the coverglass is immersed 30 min at 4°C in Lysis Solution (2.5 M NaCl, 100 mM EDTA, 10 mM Tris–HCl,1% [wt/vol] sodium lauryl sarcosine, 1% [vol/vol] Triton X-100, pH 10) and then immersed 20 min at room temperature in Unwinding Solution (200 mM NaOH, 1 mM EDTA pH 10). Unwinding is the commercial term used by the manufacturer to denote DNA denaturation. Electrophoresis is then run at 21V 30 min in Alkaline Electrophoresis Buffer (200 mM NaOH,1 mM EDTA pH8) in a CometAssay ® Electrophoresis System. Slides are then rinsed 2 × 5 min with H_2_O and 1 × 5 min with 70% EtOH, dried 30 min at 37°C, DNA dyed with SYBR ® Gold 1:10,000 and neutral lipids with AUTODOT^TM^ at 1:100, both at the same time, for 30 min, rinsed with H_2_O, and completely dried at 37°C.

### Comet Acetone

The classical comet assay is divided in 3 big steps: Lysis (1), Denaturing (2) and Electrophoresis (3). For the experiments presented in Figure 2, an additional step is intercalated at either each of these steps, whereby the coverglass is incubated at - 20°C in 100% cold acetone for 30 min. For experiments of Figure 3, steps (1) and (2) were fully replaced by incubation at -20°C in 100% cold acetone for 30min.

**Figure 2.**
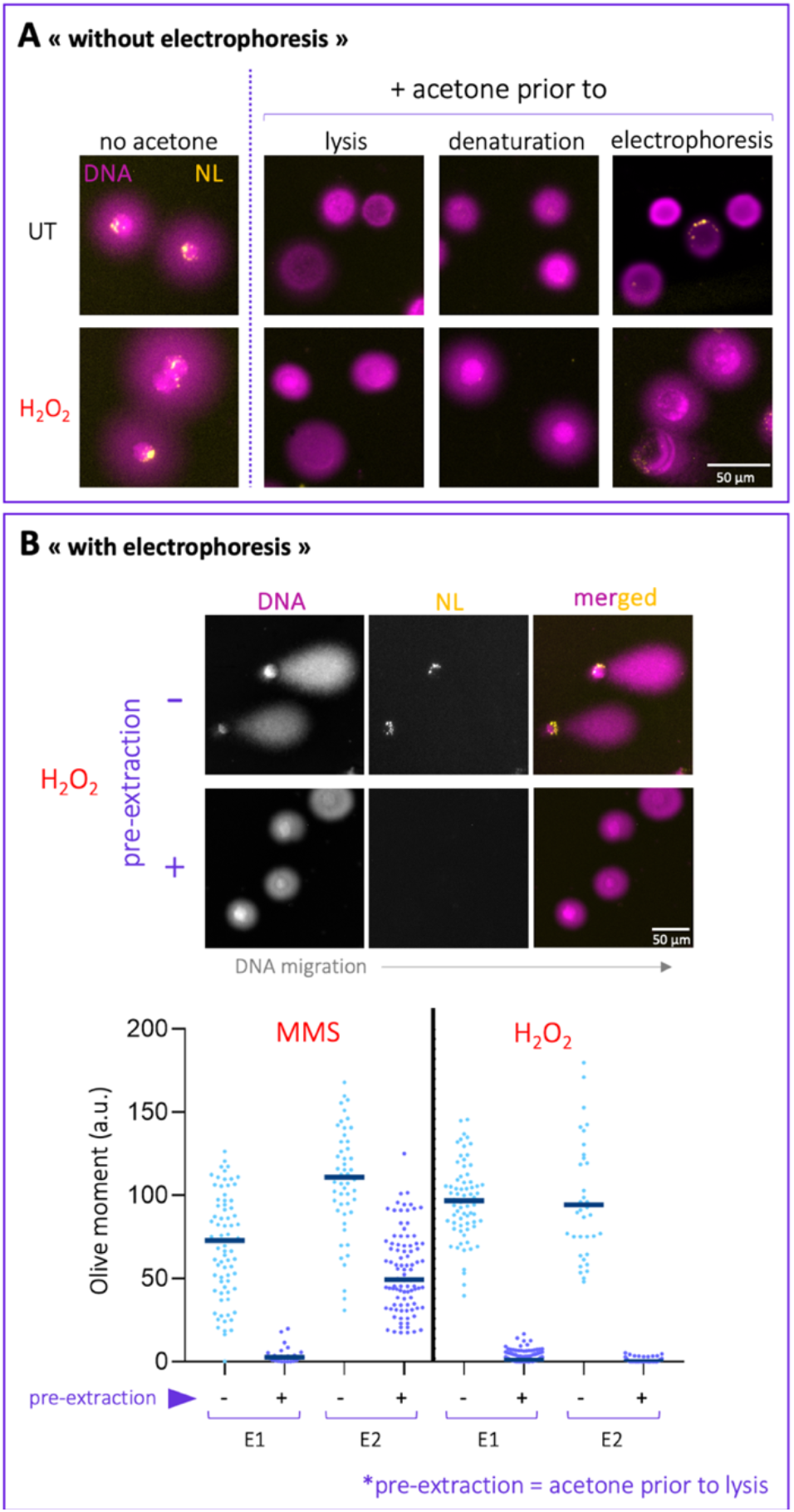
Acetone effectively removes NL from comet samples. **A**. Acetone incubation of the comet samples was performed before each of the three main steps of the comet assay protocol (prior to lysis, or to denaturation, or to electrophoresis). Importantly, electrophoresis was not accomplished in this experimental dataset. Samples were prepared from untreated harvested cells that were exposed in the collection tube to H_2_O_2_ to break DNA, or not (UT). **B. Top:** Cells treated as in (A) were used but, this time, the samples were pre-extracted using acetone prior to lysis (or not) then subjected to electrophoresis to reveal the migration of broken DNA molecules. **Bottom:** Four independent experiments were performed to break DNA either *in cellulo* (using MMS, 2 experiments) or *in vitro* (adding H_2_O_2_ to the collected cells, 2 experiments), then pre-extraction with acetone done or not. The graph represents the Olive moment measured for individual comets (dots) belonging to each experiment.

**Figure 3.**
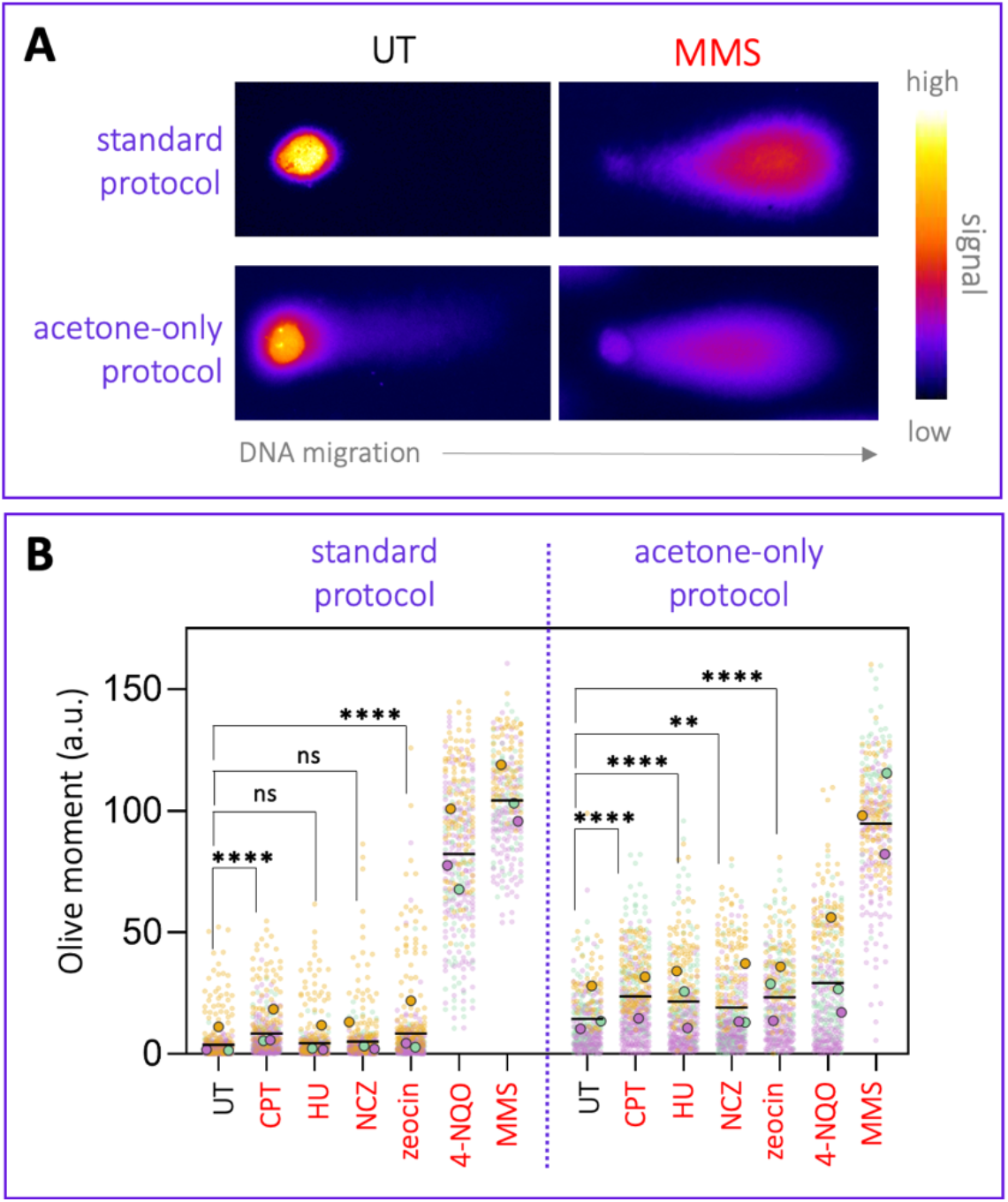
Acetone alone recapitulates the alkaline comet assay steps. **A**. The simple substitution of lysis + denaturation by acetone incubation permits the release of broken DNA molecules (cells had been exposed to MMS, or not (UT)). **B**. The performance of the acetone-only protocol (right) was compared to the standard one (left) in response to a palette of genotoxins classically used in the laboratory to create DNA damage (in red). We measured both the Olive moment, a readout of both quantity of broken DNA and the relative size of the broken fragments. Each small dot is an individual comet event. Each color is an independent experiment. Bigger dots represent the mean of each independent experiment. The horizontal black line is the mean of those means. A one-way ANOVA was applied to compare the effect of four genotoxins *versus* the untreated condition for each kind of protocol. **, *p* < 0.01; ****, *p* < 0.0001; ns, non-significant.

### Imaging and analysis

Comets and lipids were visualized under an upright microscope Zeiss Axioimager Z2 controlled either by ZEN or by Metamorph softwares, at x10 magnification. x63 magnification was only used once to have more resolutive pictures of neutral lipids. Measure of nuclei and nucleoid signals’ area, was done with Image J. Analysis of comets is performed in a semiautomated manner, using TriTek CometScore 2.0.0.38 software. Between 50 and 200 comets were analyzed per condition and experiment, and the resulting Olive moment, plotted. The Olive moment is a datum expressed in arbitrary units, resulting from the multiplication between the percentage of DNA in the tail and the distance between the centers of mass of the comet’s head and the tail. It is expressed by the following formula, where (Y_H_, X_H_) and (Y_T_ and X_T_) correspond to the coordinates of each center of mass, head and tail, respectively; and D_H_ and D_T_ correspond to the DNA in the head and in the tail (sum of gray values), respectively:

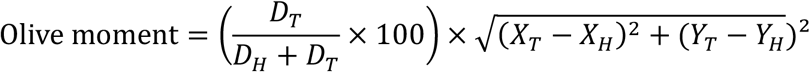

### Neutral lipid staining on cells

Cells were fixed on coverslips with 4%PFA/1X PBS for 20min, washed once with PBS and permeabilized with 0.2% Triton/1X PBS for 10 min at room temperature and then blocked with 3% BSA/1X PBS for 30 min. Coverslips were incubated with 5 μg/mL NileRed/1XPBS for 1 hour, then washed 3 times with 1XPBS under gentle shaking, then incubated for 10 min with 1 μg/mL DAPI/H_2_O, and washed 3 times with H_2_O. Finally, coverslips were allowed to dry and mounted using ProLong and then left to dry overnight at room temperature in the darkness.

### Plots and statistical analysis

Plots were made with GraphPad Prism. In scatter point graphs, each dot corresponds to one comet event and, when there is more than one color, each color corresponds to one independent repeat of the experiment. Bigger dots of different colors correspond to the means of each experiment, and black horizontal lines correspond to the mean of those means, altogether constituting Superplots (Lord *et al*, 2020). Statistical analyses were chosen accordingly to the needs and such choices are justified in the corresponding figure legends.

## Results

### Neutral lipids become trapped in the nucleoid during the comet assay protocol

During the execution of a comet protocol in human cells, paradigmatically in immortalized retinal pigmented epithelium (RPE-1) ones, fluorescence microscopy can be used at each step to monitor the space occupied by DNA signals, as revealed with SYBR™ Gold. The pressure exerted onto cells’ nuclei by the agarose pad in the first step of the protocol already increases the DNA signals’ size (Figure 1B, in pad). Further, likely as a consequence of the removal of the nuclear envelope during lysis, which removes a confinement, the signals further expand (Figure 1B, L). Later on, DNA denaturation, which separates both DNA strands, leads to a further deployment of DNA signals (Figure 1B, L+D). In these two late steps of the protocol, the DNA appears in two phases of different intensity: a signal-dense core, and a signal-light halo (Figure 1B). These structures, which can no longer be called a nucleus, are termed nucleoids.

Frequently, we detected “black holes” within the in-pad nuclei and, more markedly, within the nucleoids. These holes define in-DNA regions devoid of DNA dye, suggesting that another type of molecule was harbored within (Figure 1C, left). Of note, this observation has been informally reported in the net by other researchers (https://www.researchgate.net/post/COMET_assay_in_neutral_condition_Why_do_I_observe_black_dot_in_nucleus), although these observations have never been pursued further. These holes were visually reminiscent to nuclear lipid droplets (nLD), specific organelles that are born from the inner nuclear membrane into the nucleoplasm and which harbor neutral lipids (NL) (Sołtysik *et al*, 2021), (Figure 1C, right). While it was unlikely that these structures were actual nLD (otherwise we would systematically detect them in the original cells), it could be that the steps needed to accomplish the protocol artificially pushed within the nucleoids NL from other cellular locations, likely the cytoplasm (Figure 1D, “cell”). To test this, we incubated our samples with a dye labeling NL, AUTODOT^TM^. The in-pad samples demonstrated clustering of AUTODOT^TM^-positive signals within the DNA ones (Figure 1D). The lysed (L) and denatured (D) samples demonstrated a perfect colocalization of discrete NL signals at in-DNA holes (Figure 1D). Migration of samples arising from cells treated with the DNA-damaging agent methyl methanosulfonate, MMS, which harms DNA thus provokes the migration of negatively charged broken DNA out of the nucleoid region (*i*.*e*. tails), did not lead the AUTODOT^TM^-positive signals to migrate, suggesting that indeed they lack charge, compatible with them being NL (Figure 1E, MMS). To reinforce the notion that these NL arise from cytoplasmic NL stores, presumably cytoplasmic lipid droplets, we incubated the cells with oleate, which boosts its storage within this type of structures (Lagrutta *et al*, 2017), and re-assessed the potential accumulation of NL within comet DNA signals. We could observe a marked accumulation of NL within the DNA nucleoids (Figure 1E, oleate), confirming this interpretation. Importantly, this was not an artifact due to the dyes used to visualize DNA or lipids, as alternatively using DAPI and BODIPY to dye these molecules, respectively, did not alter the observation (Figure 1F). Thus, we conclude that during the protocol to perform comet assays, the NL present in the cytoplasm of the original cells cluster within the nucleoids.

### The neutral lipids within nucleoids can be extracted by acetone

We then wanted to assess if the presence of these NL molecules within the nucleoids altered the migration of broken DNA molecules, which could impact the conclusions reached when using this technique to assess the level of broken DNA in experimental samples. We chose to use acetone, for it has been firmly reported as a very efficient de-lipidating agent (DiDonato & Brasaemle, 2003). We incubated our samples with acetone at different stages of the protocol, *i*.*e*. prior to lysis, denaturation or electrophoresis. In all cases, acetone worked very efficiently, removing virtually all NL signals (Figure 2A). This was true for samples in which the DNA was intact (untreated), or exposed to a treatment to damage it (H_2_O_2_) (Figure 2A).

Concerning DNA signals, upon visual inspection, acetone treatment slightly altered them. In particular, applying acetone after denaturation and prior to electrophoresis led to aberrant profiles in which the core + halo DNA aspect was lost (Figure 2A). Thus, for subsequent experiments, we chose to apply acetone prior to lysis, as we estimated this preserved better the aspect of DNA signals.

We were able at this stage to evaluate whether NL presence altered, by any means, the amount of DNA signals in the tail (DNA quantity) and / or the size of such molecules (longer tails indicate smaller DNA fragments). Unexpectedly, acetone-mediated NL pre-extraction fully prevented DNA from migrating out of the nucleoid, irrespective of whether DNA had been broken *in cellulo* (cells treated with the genotoxic agent MMS) or *in vitro* (by addition of H_2_O_2_) (Figure 2B). Thus, although acetone represents a very efficient *in vitro* treatment to remove NL from the samples, a chemical reaction likely exerted onto DNA molecules themselves, and / or with the agarose pad, and / or with the other chemicals subsequently used during the protocol, prevents DNA from migrating during the last step of the protocol, the electrophoresis.

### Acetone alone permits the implementation of a simplified comet assay protocol

Despite this limitation, during our analyses to test at which step of the protocol acetone could be best used, we realized that the simple fact of adding acetone to the in-pad samples recapitulated the whole protocol. Indeed, acetone alone substituted for all the classical steps needed to “release” DNA (*i*.*e*., lysis and denaturation) (Figure 3A). This surprising observation opened to two advantages: *1)* it is possible to enormously simplify the comet assay protocol; *2)* it allows to get rid of NL.

We therefore moved to assess how DNA breaks, as created by exposing cells to a palette of genotoxins, were monitored using the classical comet assay protocol *versus* the acetone-only one. We measured the Olive moment, a parameter that considers both the amount of DNA in the tail and the relative length of the fragments in it (Figure 3B and see Material and Methods). Our first observation was that the acetone-based protocol permitted broken molecules to leave the nucleoid area and thus allowed the detection of broken DNA (Figure 3A,B). Second, with the exception of the samples in which cells were exposed to the drugs MMS and 4-NQO, for which the protocol using acetone led to modestly lower, and drastically lower values than the classical protocol, respectively, the broken molecules created by all the remaining treatments were detected more readily using the acetone protocol than the classical one (Figure 3B, compare both plots distribution and statistical significance of the difference between means). MMS and 4-NQO provoke the chemical modification of DNA bases, in contrast to the rest, suggesting that acetone may interact with these alterations and restrict subsequent DNA migration. Bearing this in mind, the results suggest that, when using genotoxins that do not alter the chemical nature of the DNA molecule, the one-step acetone protocol: 1) provides a faster execution; 2) enhances the detection of mildly broken DNA (*i*.*e*. big yet broken), which otherwise appears almost indistinguishable from the untreated condition (Figure 3B, compare standard *versus* acetone-only for CPT, HU, NCZ, zeocin). However, an increased visualization of broken DNA was also true for the untreated condition. This could be interpreted in two ways: *a)* the acetone-only protocol harms DNA; *b)* NL in the nucleoid restrict the free migration of broken DNA molecules of big size and, in their absence, very big fragments corresponding to poorly broken DNA may become detectable even in basal conditions.

### Neutral lipids within the nucleoid restrict the migration of broken DNA molecules

To discriminate between these possibilities, we moved back to a classical protocol (no acetone) and very efficiently removed NL *in vivo* by starving the cells (Figure 4A, left, NL in cells). We performed comet assays onto the DNA of these starved cells (not exposed to any genotoxin) and asked if, in those conditions, the DNA, free of NL, could migrate further than in the non-starved condition. We observed that this was not the case, in contrast to the increase in Olive moment of untreated cells subjected to the acetone-only protocol (Figure 4B). This experiment suggests that performing the comet assay using an acetone-only protocol leads to mild damage of unbroken DNA molecules that manifests as an increase in fragments of long size in basal measurements (Figure 3B & 4B).

**Figure 4.**
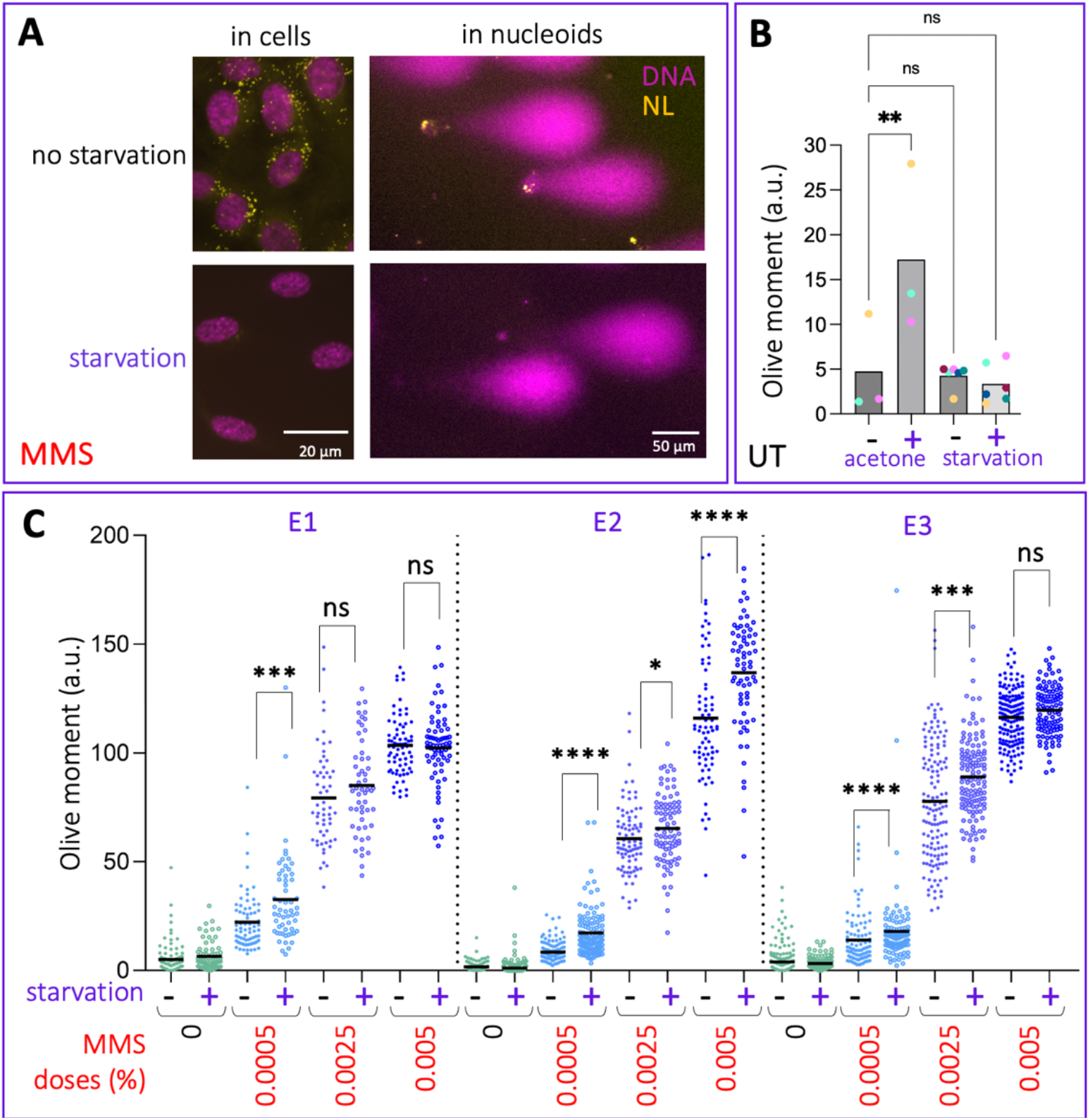
Acetone and NL impact the migration of DNA during comet assays. **A**. Cells were starved (or not) to remove NL (left) then exposed to MMS to break DNA and standard comet assay was performed (right). **B**. Graph comparing the mean Olive moment obtained from untreated cell samples that were processed either using the standard protocol (-), or prepared using the acetone protocol (+ acetone); or the standard protocol but using cells previously starved (+ starvation). The colors refer to independent experiments. The data shown for acetone (-, +) are taken from Figure 3B. The data used for starvation (-, +) are new experiments. Each bar height shows the mean of the individual means. A one-way ANOVA, pertinent for multiple comparisons, was applied to assess if the means of the means were significantly different. **, *p* < 0.01; ns, non-significant. **C**. Three independent experiments (E1, E2, E3) were performed to compare samples arising from non-starved *versus* starved cells that had been exposed, or not, to the indicated doses of MMS. Each dot is an individual comet event. The horizontal black line is the mean of those dots. To assess whether the populations displayed different averages, and given that the population distributions varied, non-parametric Mann-Whitney tests were applied. *, *p* < 0.05; **, *p* < 0.01; ***, *p* < 0.001; ****, *p* < 0.0001; ns, non-significant.

We also treated the starved and the non-starved cells with MMS (for which no artifacts were expected in the absence of acetone), as this drug yields very patent broken DNA-containing tails (Figure 4A, right). We used MMS in a range of doses commonly used on cultured human cells (0.0005%, 0.0025%, 0.005%). As expected, the tails from the samples after electrophoresis became increasingly longer as the MMS dose increased (Figure 4C). Importantly, NL removal was consistently accompanied by higher Olive moments than those obtained from non-starved cells at any MMS dose (Figure 4C). Despite inter-dose and inter-experimental variability, the trend was perfectly respected in all cases and led to highly significant differences in most of them (Figure 4C). These data imply that NL in the nucleoid restrict the migration of broken DNA molecules into the tail during standard comet assays.

## Discussion

In this work, we have demonstrated that NL present in cells become trapped within the DNA mass (the nucleoid) during the preparation of the samples for comet assay. These lipids later on interfere with the ability of the broken DNA molecules, present within the nucleoid, to migrate out of it and into the tail, whose detection is the ultimate goal of performing comet assays. De-lipidation *in vivo*, for example by starvation, solves this concern, however this intervention may drastically affect the physiology of the cells under investigation. Alternatively, we show that NL can be removed *in vitro* by performing a simpler comet assay with acetone, although this manipulation seems to create a modest DNA damage that manifests, in untreated samples, as slowly migrating (big) DNA fragments. Last, we uncover a limitation of this acetone-based protocol when the genotoxin under consideration modifies the chemistry of the DNA molecule.

The reason why NL become trapped within the DNA mass is very likely due to the expansion suffered by the DNA during the implementation of the protocol, as already trapping the cells under the agarose pad increases the DNA-occupied area (Figure 1B). Later on, nuclear envelope disintegration by lysis further makes the DNA extend onto previously-in-the-cytoplasm NL (Figure 1D). Our data suggest that, once trapped, the NL interact with the DNA molecules and modify their migration ability when exposed to an electric field.

Our systematic assessment of the ability of acetone to extract NL from comet assay samples led us to realize that acetone alone is a simple, fast and reliable manner to simplify the comet assay protocol while getting rid of NL. This is likely due to the fact that the extraction capacity of acetone (DiDonato & Brasaemle, 2003) may not be restricted to lipids, thus may help remove other cellular components, thus bypassing the need to do lysis and unwinding. Moreover, the differences seen between the untreated and the genotoxin-treated samples, which for some of these agents is frequently very modest, become more marked (Figure 3B, CPT, HU, NCZ, zeocin) when using the acetone-only protocol. This is a neat advantage when monitoring subtle increases in DNA breaks, as is the case for some treatments or experimental conditions. However, two limitations apply to this tool: first, some genotoxins modify the chemical nature of DNA, such as the alkylating drugs MMS and 4-NQO (Prakash & Strauss, 1970; Mirzayans *et al*, 1985; Nagao & Sugimura, 1976). Acetone is likely to alter the interaction between this modified DNA and the NL, and / or to create artifactual connections with the agarose matrix. Thus, before the simplified acetone protocol is to be used in a regular manner, the genotoxin under consideration must be first compared using the standard and the acetone-only protocol. Second, we detected that in untreated (thus, likely undamaged DNA) samples, the acetone-only protocol leads to detection of broken DNA molecules (Figure 3B). These molecules are little damaged, *i*.*e*. migrate close to the nucleoid (Figure 3A) and as such display low Olive moments (Figure 3B), yet cannot be attributed to basal damage that would be invisible to the standard method. Indeed, removal of NL by other means than acetone in undamaged cells, for example by starvation, did not lead to the same outcome (Figure 4B), suggesting that acetone alone slightly harms DNA.

Last, we observed that broken DNA molecules existing prior to the comet assay protocol implementation are more easily detected if NL are removed, as shown when using starvation (Figure 4C, MMS). The unsaturated fatty acid oleate, as the one used in Figure 1E, and frequently used in the literature to trigger NL storage within cytoplasmic reservoirs, does so by becoming esterified into triacylglycerol (TAG) molecules. Oleate has been reported to directly bind DNA *in vitro*. It recognizes A-T base pairs and can bind them tightly at a ratio of one oleate molecule per 2–3 base pairs (Zhdanov *et al*, 2002). This may underlie the reason why the lipidomic characterization of the lipids bound to DNA in *Pseudomonas aurantiaca* revealed a very tight binding of TAG to DNA (Zhdanov *et al*, 2015). This can seem surprising, given that DNA is an overall negatively charged electrolyte, while TAGs are neutral molecules. However, the fact that oleate can bind A-T base pairs suggest that NL interaction with DNA may occur in the central DNA groove, and not with its backbone. In any case, both reports are in agreement with our observations, and would suggest that NL within the DNA mass of the nucleoid, to some extent, retain broken DNA molecules, thus restricting their migration during the electrophoresis.

Beyond technical considerations, our work raises the notion that NL in direct contact with DNA, for example in the context of a cell, may exert regulatory or structural functions. As briefly mentioned, nuclear lipid droplets exist in the nucleoplasm of cells (reviewed in (Samardak *et al*, 2024)) and they harbor NL (Layerenza *et al*, 2013). The secretion of LD contents with chemical chaperone activities is described (Moldavski *et al*, 2015), and alterations on chromatin behavior are dependent on NL delivered by nuclear lipid droplets (Umaru *et al*, 2023). Our report adds, to the frame of nuclear physiology, the possibility of a control exerted by NL by locally restricting DNA mobility.

## Abbreviations

MMS: methyl methanesulfonate
NL: neutral lipids
TAG: triacylglycerol.

## Author contributions

Conceptualization, C.S., J.J.-C. and M.M.-C.; methodology, C.S. and M.M.-C.; validation, C.S.; formal analysis, C.S., J.J.-C. and M.M.-C.; investigation, C.S. and J.J.-C.; writing—original draft preparation, C.S. and M.M.-C.; writing—review and editing, C.S., J.J.-C. and M.M.-C.; visualization, C.S. and M.M.-C.; supervision, M.M.-C.; project administration, M.M.-C.; funding acquisition, M.M.-C. All authors have read and agreed to the published version of the manuscript.

## Acknowledgements

We thank the imaging facility MRI, a member of the national infrastructure France-BioImaging, supported by the French National Research Agency (ANR-10-INBS-04, Investissements d’avenir). We thank the French National Research Agency for supporting the execution of this work (ANR-21-CE12-0004-01).

## Competing interests

The authors declare no competing interests

## Notes

### Competing Interest Statement

The authors have declared no competing interest.

